# wQFM-DISCO: DISCO-enabled wQFM improves phylogenomic analyses despite the presence of paralogs

**DOI:** 10.1101/2023.12.05.570122

**Authors:** Sheikh Azizul Hakim, MD Rownok Zahan Ratul, Md. Shamsuzzoha Bayzid

**Affiliations:** Department of Computer Science and Engineering, Bangladesh University of Engineering and Technology Dhaka-1205, Bangladesh

**Keywords:** Species tree, gene tree, gene tree heterogeneity, incomplete lineage sorting (ILS), Gene duplication and loss (GDL), multicopy gene trees, Paralogy

## Abstract

Gene trees often differ from the species trees that contain them due to various factors, including incomplete lineage sorting (ILS), gene duplication and loss (GDL), and horizontal gene transfer (HGT). Several highly accurate species tree estimation methods have been introduced to explicitly address ILS, including ASTRAL, a widely used statistically consistent method, and wQFM, a quartet amalgamation approach that is experimentally shown to be more accurate than ASTRAL. Two recent advancements, ASTRAL-Pro and DISCO, have emerged in the field of phylogenomics to consider gene duplication and loss (GDL) events. ASTRAL-Pro introduces a refined measure of quartet similarity, accounting for both orthology and paralogy. DISCO, on the other hand, offers a general strategy to decompose multicopy gene family trees into a collection of single-copy trees, allowing the utilization of methods previously designed for species tree inference in the context of single-copy gene trees. In this study, we first introduce some variants of DISCO to examine its underlying hypotheses and present analytical results on the statistical guarantees of DISCO. In particular, we introduce DISCO-R, a variant of DISCO with a refined and improved pruning strategy that provides more accurate and robust results. We then propose wQFM-DISCO (wQFM paired with DISCO) as an adaptation of wQFM to handle multicopy gene trees resulting from GDL events. Extensive evaluation studies on a collection of simulated and real data sets demonstrate that wQFM-DISCO is significantly more accurate than ASTRAL-Pro and other competing methods.

## 1 Introduction

Inferring species trees from gene trees sampled throughout the whole genome is a fundamental problem in molecular evolutionary biology. However, this task is complicated by the phenomenon of gene tree heterogeneity (or discordance), where different genes may have different evolutionary histories. Gene tree heterogeneity can arise due to various biological processes, including incomplete lineage sorting (ILS), gene duplication and loss (GDL), and horizontal gene transfer (HGT) (Warnow, 2017). Numerous methods have been proposed for species tree inference, including co-estimation of gene trees and the species tree (Boussau et al., 2013a; Heled and Drummond, 2010; Liu, 2008; Szöllosi et al., 2015), and species tree inference from sequence data (Bryant et al., 2012; Chifman and Kubatko, 2014). However, the most scalable and popular approach to date remains a two-step process, where gene trees are first inferred independently from sequence data, and then combined using summary methods that attempt to estimate species trees by summarizing the input gene trees under a model of gene tree discordance.

Many summary methods have been proposed to target ILS and some of them have been shown to be statistically consistent under the multi-species coalescent (MSC) model (Avni et al., 2015; Bayzid and Warnow, 2012; Chifman and Kubatko, 2014; Islam et al., 2020; Mahbub et al., 2021; Mim et al., 2023; Mirarab et al., 2014; Reaz et al., 2014; Snir and Rao, 2010; Zhang, 2011). Other statistically consistent species-tree estimation methods include BEST (Liu, 2008) and *BEAST (Heled and Drummond, 2010), which co-estimate gene trees and species trees from input sequence alignments. While these methods can produce substantially more accurate trees than other methods, they are extremely computationally intensive and do not scale to large numbers of genes (Bayzid and Warnow, 2012, 2013; Smith et al., 2013).

Maximizing quartet consistency (MQC) is one of the leading optimization criteria for estimating statistically consistent species trees from gene trees in the presence of ILS (Avni et al., 2015; Mahbub et al., 2021; Mirarab et al., 2014; Reaz et al., 2014; Snir and Rao, 2010). MQC seeks a species tree that is consistent with the largest number of quartets induced by the set of gene trees. ASTRAL is arguably the most commonly used quartet-based summary method (Mirarab et al., 2014). Given a set of unrooted gene trees, ASTRAL uses a dynamic programming approach to solve the *NP* -hard optimization problem of finding the species tree that agrees with the largest number of quartet trees induced by the set of gene trees. A different method involves deducing individual quartets and then combining them into a cohesive species tree in a divide-and-conquer fashion, either with or without weights, e.g., Quartets Max-Cut (QMC) (Snir and Rao, 2010), Quartets Fiduccia–Mattheyses (QFM) (Reaz et al., 2014), Weighted Quartets Max-Cut (wQMC) (Avni et al., 2015), Weighted Quartets Fiduccia–Mattheyses (wQFM) (Mahbub et al., 2021) etc. While ASTRAL is an exact dynamic programming algorithm statistically consistent under the MSC model, wQMC and wQFM are heuristic-based methods and no result on statistical consistency exists in the literature. However, through extensive simulation studies, wQFM has been shown to have consistently outperformed wQMC and ASTRAL (Mahbub et al., 2021, 2022), even when gene trees are incomplete (Mahbub et al., 2022). These methods were originally proposed to address gene tree heterogeneity due to ILS and are thus applicable to single-copy gene trees.

GDL events introduce paralogs, which are genes originating from a common ancestor through duplication events, in addition to orthologs, which arise from speciation events (Fitch, 2000; Yang and Smith, 2014). Consequently, gene families may consist of multiple genes with the same species labels, resulting in multicopy gene trees. However, most of the summary methods require genes with at most one copy in each species, and so cannot be used directly in the presence of GDL. One way to address this is to identify the orthologous genes so that the existing summary methods that were designed to handle single-copy gene trees can be applied to estimate species trees from sets of orthologous genes. However, reliable identification of orthologous genes is challenging (Altenhoff et al., 2019). Furthermore, relying solely on orthologous genes leads to the exclusion of numerous multicopy gene trees, thereby discarding valuable information (1kp, 2019; Wickett et al., 2014).

There have been a few summary methods developed to estimate species trees from multicopy gene family trees by addressing gene duplications and losses and without requiring orthology determination, namely PHYLDOG (Boussau et al., 2013a), iGTP (Chaudhary et al., 2010), DupTree (Wehe et al., 2008), DynaDup (Bayzid, 2023; Bayzid and Warnow, 2018; Bayzid et al., 2013), MulRF (Chaudhary et al., 2013), FastMulRFS (Molloy and Warnow, 2020), Guenomu, ASTRAL-Pro (Zhang et al., 2020), and Species-Rax (Morel et al., 2022). Most of these methods (Bayzid and Warnow, 2018; Bayzid et al., 2013; Chaudhary et al., 2010, 2013; Wehe et al., 2008) are parsimony-based approaches where a species tree is estimated by minimizing a particular distance between the species tree and given set of gene trees (e.g., the total number of duplication and loss events). However, there are other methods that are more agnostic about the reasons for gene tree discordance and do not necessarily rely on maximum parsimony reconciliation. Example methods of this type include PHYLDOG (Boussau et al., 2013a) and Guenomu (De Oliveira Martins et al., 2016) that represent a powerful approach, leveraging probabilistic models and Bayesian techniques, which co-estimates gene trees and species trees. These methods are computationally very demanding, so they are limited to small datasets. ASTRAL-Pro is a quartet-based species-tree inference method that aims to compute a species tree that maximizes the total similarity to the input gene trees, using a new measure of quartet similarity that accounts for orthology and paralogy (Zhang et al., 2020).

Another approach to tackle multicopy gene family trees resulting from GDL events is to decompose multicopy gene trees into single-copy gene trees (Dunn et al., 2013; Hejnol et al., 2009; Kocot et al., 2013). Willson et al. (2021) introduced the DISCO algorithm to decompose multicopy gene trees into single-copy gene trees, allowing the utilization of methods designed for single-copy gene trees in handling multicopy gene trees. To obtain a rooted tagged (duplication/speciation) multi-labeled tree, DISCO utilizes the tagging and rooting heuristics of ASTRAL-Pro and uses a pruning strategy to produce single-copy gene tress from multicopy gene family trees. ASTRAL-Pro and ASTRAL-DISCO (ASTRAL on the output set of DISCO) have been shown to be statistically consistent when either ILS or GDL is active (Willson et al., 2021; Zhang et al., 2020). Many successful summary methods (ASTRAL, wQFM, MP-EST, etc.) were designed to model ILS and therefore, suitable for only single-copy gene trees. Thus, any fundamental improvements of DISCO and its applications to enhance the capabilities of highly accurate methods such as wQFM in modeling both orthology and paralogy, thereby making them suitable for multicopy gene trees will be of great interest for estimating species trees from multi-locus data in the presence of GDL.

This study entails the following key contributions. We first examine some underlying hypotheses regarding the pruning and tagging strategies of DISCO combined with both ASTRAL and wQFM and proposed DISCO-R (DISCO with a **R**efined pruning strategy that **R**egrafts clades that original DISCO may have missed), which resulted in more accurate and robust results. We for the first time apply wQFM in the multicopy gene tree setting and establish its experimental superiority over other competing methods including ASTRAL-Pro. Our findings from an extensive evaluation study highlight the effectiveness of the combined wQFM-DISCO approach and emphasize its superiority in species tree estimation from multicopy gene trees. We also extend theoretical guarantees of statistical consistency of ASTRAL combined with DISCO by showing that ASTRAL-DISCO is statistically consistent under DLCoal (Duplication, Loss, and Coalescence) model – a model to incorporate both ILS and GDL (Rasmussen and Kellis, 2012).

## 2 Background and preliminaries

We now give a brief overview of the DLCoal model, ASTRAL-Pro, DISCO, and wQFM to make this paper self-contained.

### 2.1 DLCoal model

The following brief overview of the unified duplication-loss-coalescence (DLCoal) model has been adapted from (Markin and Eulenstein, 2021; Rasmussen and Kellis, 2012).

A *species tree* represents an evolutionary history of a group of species. A *locus tree* represents a duplication-loss history of a particular gene. A locus tree is obtained from a species tree by running the duplication/loss process top-down along the edges of the species tree. In other words, the duplication/loss process is a birth-death process with a fixed birth (duplication) rate and death (loss) rate (Arvestad et al., 2004). The birth-death process starts in the root edge of the species tree; whenever it reaches a speciation point, the process splits into two copies and continues independently in the children edges. A locus tree node can be termed as *speciation* if such a node corresponds to a speciation event/node in the species tree or as *duplication* if such a node corresponds to the creation of a new locus. Note that locus tree leaves are labeled by gene names. The *gene tree* is obtained from a locus tree by running the bounded multispecies coalescent (b-MSC) process bottom-up along the edges of the locus tree. The duplication/loss process and the b-MSC process are jointly referred as the DLCoal model. In the standard multispecies coalescent (MSC) model, there is exactly one gene lineage starting in every extant locus tree leaf, and if two or more lineages enter the same locus tree edge, then the coalescence history of these lineages is determined by an exponential distribution. The b-MSC process in the unified DLCoal model is defined by imposing constraints on MSC due to the duplication points. In particular, all lineages originating below a daughter duplicate must coalesce below the respective duplication node.

### 2.2 ASTRAL-Pro

We begin this section by reviewing the terminology introduced in (Zhang et al., 2020). Let *S* be a set of *n* species. The input to our problem is a collection of multicopy gene trees. Given a rooted multicopy gene tree *G*, we define a many-to-one mapping *α*_*G*_ between its leaf-set ℒ_*G*_ and *S*. For a leaf *l* ∈ℒ_*G*_, *α*_*G*_(*l*) denotes the species the gene *l* was taken from. For a node *u*, we define ℒ_*G*_(*u*) to be the set of leaves under *u*. We extend definitions such that *α*_*G*_(*A*) = *{α*_*G*_(*a*)|*a ∈ A}* for *A ⊂* ℒ_*G*_ and *α*_*G*_(*u*) = *α*_*G*_(ℒ_*G*_(*u*)) for a node *u* in *G*. In other words, *α*_*G*_(*u*) denotes the set of species whose genes are present under a node *u* in *G*.

A quartet *Q* is a four-membered subset ℒ_*G*_. If we recursively delete leaves from *G* such that only four leaves in *Q* are left, we are left with a tree with two degree-3 nodes. These two nodes are called the anchors of *Q*. Since *G* is rooted, we can find the LCA of these two nodes which is referred to as the anchor LCA of *Q* in *G*.

We tag every non-leaf node as speciation or duplication. Given a non-leaf node *u* with two children *u*_1_ and *u*_2_, we tag *u* as a speciation node if *α*_*G*_(*u*_1_) *∩ α*_*G*_(*u*_2_) = *{}*. Otherwise, we tag *u* as a duplication node. A quartet with a topology *ab*|*cd* in a gene tree is defined as a speciation-driven quartet (SQ) if all the four genes are contained in different species and the LCA of either *a* or *b* with either *c* or *d* is tagged as speciation. Two SQs on the same four species are equivalent if they have the same anchor LCA. The per-locus quartet score of a species tree S with respect to a rooted tagged gene tree G is the number of equivalent quartet classes that match the S topology. The *Maximum per-Locus Quartet score Species Tree* (MLQST) problem is to find the species tree that maximizes the PL quartet score with respect to input gene trees, given a set of rooted tagged gene trees. ASTRAL-Pro, in its exact version, solves the MLQST problem.

ASTRAL, on the other hand, was designed to solve the *Maximum Quartet Support Species Tree* (MQSST) problem, which aims to find the species tree that agrees with the largest number of quartet trees induced by the set of gene trees. ASTRAL, or in general, the solution to the MQSST problem, is statistically consistent under MSC (Mirarab et al., 2014).

### 2.3 DISCO

ASTRAL-Pro roots and tags using the maximum parsimony principle with equal penalties for duplication and loss, and no penalty for ILS. DISCO performs a post-order traversal of the multicopy gene tree and prunes the smaller subtree at each duplication node.

The central idea of DISCO is to decompose each multicopy gene tree into a set of single-copy gene trees (Willson et al., 2021). To accomplish so, DISCO uses the rooting and tagging algorithm of ASTRAL-Pro. ASTRAL-Pro roots and tags using the maximum parsimony principle with equal penalties for duplication and loss, and no penalty for ILS. Given a fully resolved unrooted binary tree, the ASTRAL-Pro algorithm places the root along an edge such that the total number of duplications and losses is minimized. After rooting the tree, each internal node is tagged either as a duplication or a speciation node. Given a rooted tagged gene tree *G*, the set of output trees *S*_*G*_ is initially empty. The decomposition algorithm performs a postorder traversal. When it reaches a duplication node, the smaller subtree rooted at that node is pruned and added to *S*_*G*_. When the entire traversal is complete, the resultant tree is also added to *S*_*G*_. It can be ensured that every tree in *S*_*G*_ is a single-copy tree. Summary methods take a collection of gene trees *C* = *{G*_*i*_*}* as input. The output of DISCO when fed with *C* as input is 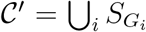 . An illustrative example is presented in Section 4.

### 2.4 wQFM

wQFM considers the weighted version of the *Maximum Quartet Consistency* (MQC) problem (Reaz et al., 2014; Snir and Rao, 2010), which we call *𝒲ℳ𝒬𝒞*, and define as follows.

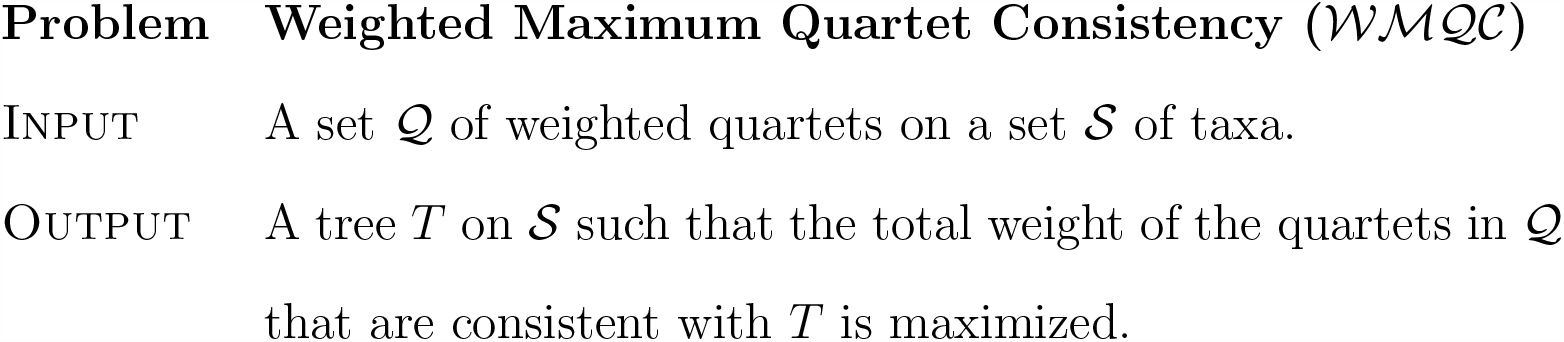

wQFM, in general, can be used to amalgamate any set of given weighted quartets (e.g., quartets induced by a the gene trees or the quartets produced by SVDquartets). In this study, we consider wQFM-GTF which estimates species trees by combining the quartets induced by the gene trees, and the weights of the quartets are computed based on the gene tree frequency (GTF), i.e., we use the frequency of the quartets in input gene trees as weights.

## 3 Statistical consistency of ASTRAL-DISCO under the unified DLCoal model

Willson et al. (2021) showed that ASTRAL-DISCO is statistically consistent when the cause of discordance is only GDL, provided that the rooting and tagging algorithm of ASTRAL-Pro is correct. Legried et al. (2020) proved the statistical consistency of ASTRAL-ONE under GDL. The ASTRAL-ONE pipeline consists of two stages. First, given a collection of multicopy gene trees, from each multicopy tree, a single-copy gene tree is obtained by randomly sampling one gene from each species. Then ASTRAL is run on the single-copy gene trees. Markin and Eulenstein (2021) extended the results of Legried et al. (2020) when ILS is also involved besides GDL. Under the unified (DLCoal) process (Rasmussen and Kellis, 2012), Markin and Eulenstein (2021) proved the following theorem.

### Theorem 1

(Markin *et al*., 2021). *Let S be a species tree with four leaves that displays quartet AB*|*CD, and let G be a gene tree that evolved in S according to the DLCoal process. If one picks genes a, b, c, d (that correspond to species A, B, C, D respectively) uniformly at random (assuming they exist) from G, then P* (*ab*|*cd ∈ G*) *> P* (*ac*|*bd ∈ G*) = *P* (*ad*|*bc ∈ G*).

When the inequality in Theorem 1 is satisfied, Legried et al. (2020) proved that ASTRAL-ONE is a statistically consistent estimator as presented in Theorem 2.

### Theorem 2

(Legried et al, 2020; rephrased). *Let S be a species tree with four leaves that displays quartet AB*|*CD, and let G be an arbitrarily large collection of gene trees over the species in S. If one picks genes a, b, c, d (that correspond to species A, B, C, D respectively) uniformly at random (assuming they exist) from any G ∈ G and the resultant quartet q in G is more likely to match the species tree configuration than the other two, i*.*e*., *P* (*q* = *ab*|*cd*) *> P* (*q* = *ac*|*bd*) = *P* (*q* = *ad*|*bc*), *then ASTRAL-ONE almost surely returns S when run with G*.

We extend the previous results to show the statistical consistency of ASTRAL-DISCO under the unified DLCoal model (Theorem 3).

### Theorem 3.

*ASTRAL-DISCO is statistically consistent under the DLCoal process, provided that the input gene trees are correctly rooted and tagged*.

*Proof*. By assumption, each gene tree is obtained by running the DLCoal process over the true species tree, and the rooting and tagging are correct. Hence each duplication node in the gene tree marks the creation of a new locus. We consider that the subtree to be kept follows the parent locus (mother duplicate), and the subtree to be pruned follows the novel locus (daughter duplicate). Therefore, each resultant DISCO tree represents the history of a particular locus, which obeys the bounded-MSC model (a part of the DLCoal process). If one picks genes *a, b, c, d* (that correspond to species *A, B, C, D* respectively, assuming they exist) from any resultant DISCO tree *G*, then by Theorem 1, *P* (*ab*|*cd ∈ G*) *> P* (*ac*|*bd ∈ G*) = *P* (*ad*|*bc ∈ G*). Since a resultant DISCO tree is single-copy, there is only one quartet *q* induced by genes *a, b, c, d*, which can be either *ab*|*cd, ac*|*bd*, or *ad*|*bc*. So, *P* (*q* = *ab*|*cd*) *> P* (*q* = *ac*|*bd*) = *P* (*q* = *ad*|*bc*). Hence, by Theorem 2, ASTRAL-ONE on the resultant DISCO trees is statistically consistent under the DLCoal process. Since DISCO trees are single-copy trees, running ASTRAL on DISCO trees and ASTRAL-ONE on DISCO trees are equivalent and the theorem follows.

Since ASTRAL, both in its exact and heuristic version, exactly solves the MQSST problem when there are adequately many error-free gene trees, all the statistical consistency results for ASTRAL will hold for any other algorithm that accurately solves MQSST. In particular, if the quartet amalgamation technique is proper and the weights of the input quartets are computed based on GTF, then the unique solution to the *WMQC* problem (the problem wQFM tries to heuristically solve) is a statistically consistent estimator of the true species tree under the MSC model (Mahbub et al., 2021). Following the same argument as above, we can conclude the following theorem.

### Theorem 4.

*If the quartet amalgamation technique is proper and the weights of the input quartets are computed based on GTF of the trees decomposed by DISCO, then the unique solution to the WMQC problem is a statistically consistent estimator of the true species tree under the DLCoal model*.

## 4 DISCO-R: A refined pruning strategy using re-grafting

In this study, we tried to investigate some variants of DISCO and empirically showed that DISCO is quite robust. In particular, we propose an improved pruning strategy and call this variant DISCO-R. Zhang *et al*. introduced the concept of speciation-driven quartets (SQ) to formulate the ASTRAL-Pro algorithm (Zhang et al., 2020). They claimed that only SQs contain information about speciation. Since DISCO uses the rooting and tagging heuristic of ASTRAL-Pro, the quartets in the resultant DISCO trees are all marked as SQs and considered by ASTRAL-Pro (Willson et al., 2021). However, DISCO does not consider all speciation-driven quartets. Here, we introduce a new pruning heuristic DISCO-R, that provably includes more SQs than DISCO. We empirically show that DISCO-R performs at least as good as or better than DISCO.

Let us consider a multicopy gene tree *G* in Figure 1a. We differentiate the instances of the same species by a numerical subscript. There is one duplication node which we mark as red. Since both subtrees have the same number of leaves, DISCO could prune any of the subtrees. Without loss of generality, let us assume that the larger subtree was pruned. When fed with *G*, DISCO produces the two trees in Figure 1b. ASTRAL-Pro takes into consideration all SQs of *G*. All quartets in the output of DISCO trees are speciation driven. However, the output misses some SQs such as *ac*_2_|*d*_2_*e, ah*|*d*_2_*e, ab*|*pe, ab*|*ph* etc. We can see two types of SQs here. In case *ab*|*pe* and *ab*|*ph*, two species were from the pruned subtree and the rest two were from the backbone tree.

**Figure 1:**
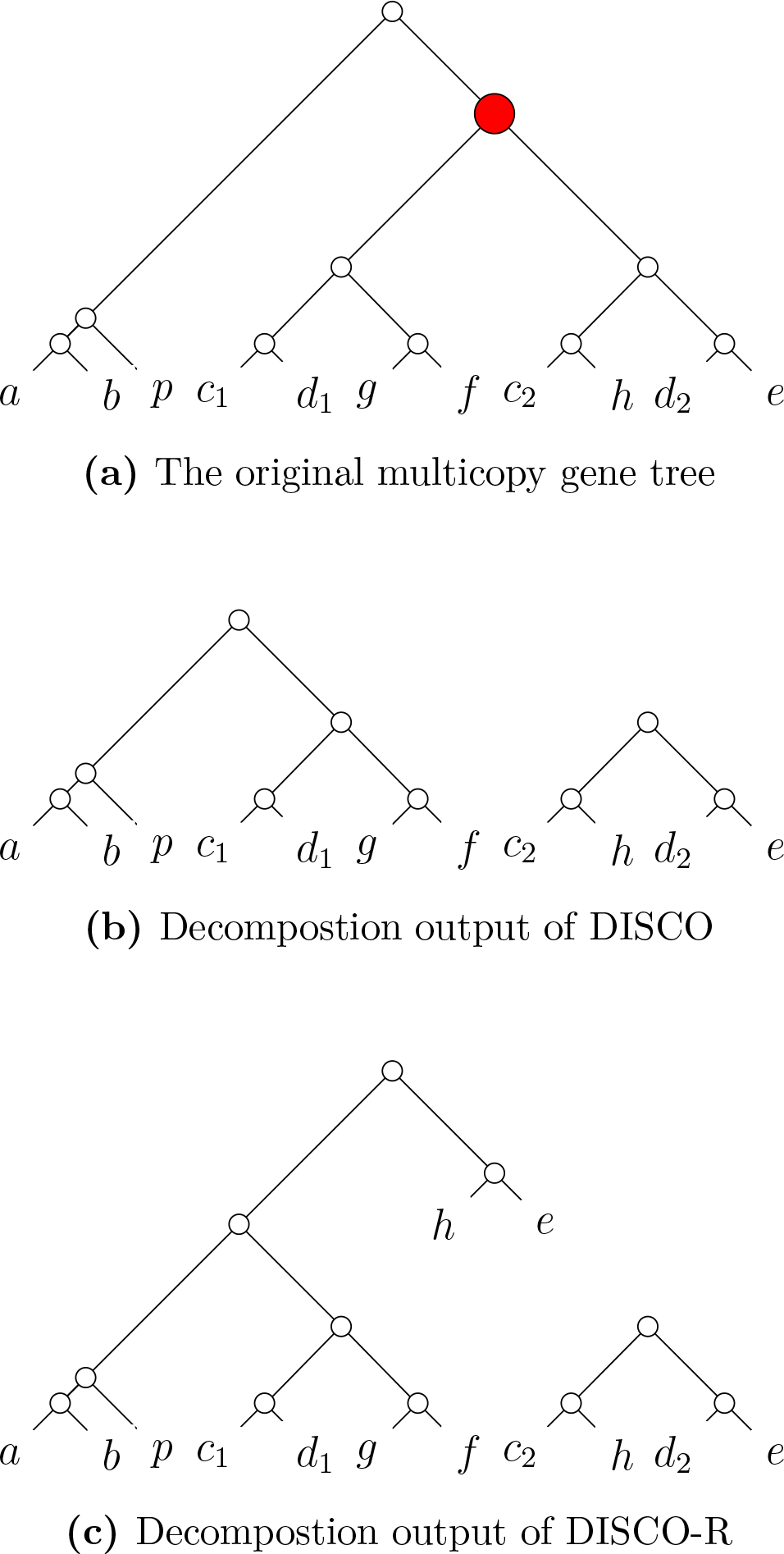
A multicopy gene tree and the decomposition outputs of DISCO and DISCO-R. The white and red nodes denote the speciation nodes and duplication nodes respectively.

We modify DISCO to include (the quartets that DISCO usually misses, ie., in our example in Fig. 1, we want to include *ab*|*pe* and *ab*|*ph*) these two quartets. In our new method DISCO-R, at a duplication node, we prune a subtree as usual. However, we take the subtree of the pruned subtree, excluding the species present in the backbone tree, and reattach it in the place of the duplication node. All degree-2 nodes are contracted. The output of DISCO-R when run with *G* is shown in Figure 1c. This scheme successfully incorporates the speciation-driven quartets *ab*|*pe* and *ab*|*ph*. However, the other SQs *ac*_2_|*d*_2_*e* and *ah*|*d*_2_*e* are still missing in DISCO-R. Moreover, DISCO-R introduces *ae*|*c*_1_*d*_1_ and *ad*_1_|*he*. These quartets are present in *G*, but they are not speciation driven per the rooting and tagging scheme of ASTRAL-Pro. In brief, the quartets generated by DISCO are SQs, but not all SQs are covered by DISCO output trees. DISCO-R might cover more SQs than DISCO, but DISCO-R can introduce some non-SQs too. ASTRAL-Pro claims that these non-SQs do not convey any speciation-related information. However, our experiments establish the efficacy of DISCO-R. Moreover, it is possible to construct a collection of gene trees *C* = *{G*_*i*_*}* such that one species is entirely missing from 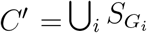 . Such events cannot occur in DISCO-R because we do not prune any non-duplicate leaf. Such an example, adapted from (Willson et al., 2021), is shown in Figure 2.

**Figure 2:**
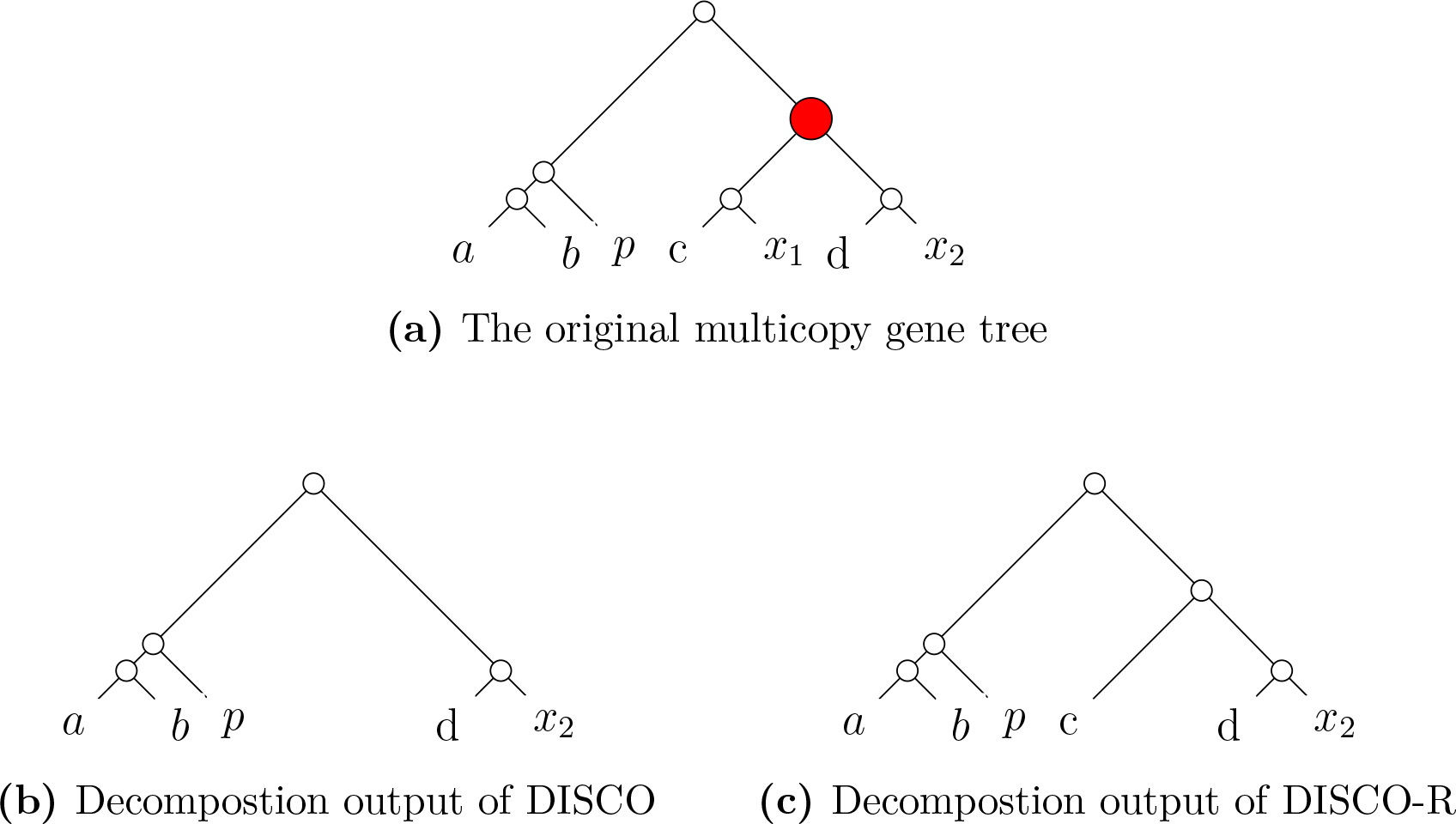
An example where DISCO misses a certain taxon (*c*) in its output. At the duplication node (marked as red), the subtree (*c, x*_1_) is pruned, which does not induce any quartet and is left out of DISCO output. However, DISCO-R regrafts the node *c* back to the gene tree

### 4.1 Variants of DISCO and DISCO-R

DISCO uses the rooting and tagging heuristic of ASTRAL-Pro where they assign equal weights to each duplication and loss event. Then, while doing a postorder traversal, the smaller subtree is pruned at each duplication node. We wanted to examine the effects of various weighting schemes and pruning strategies. We experimented with several variants of DISCO (e.g., DISCO-1, DISCO-2, …, DISCO-6). First, instead of pruning the smaller subtree at a duplication node as was done in the original DISCO algorithm, we experimented with pruning the larger subtree (DISCO-2) or randomly pruning one of the two subtrees (DISCO-3) at that node. We also experimented with the rooting algorithm of ASTRAL-Pro, where we assigned the cost of a duplication event to be twice the cost of a loss event (DISCO-5,-6). Such scoring heuristic was previously suggested in literature (David and Alm, 2010; Zhang et al., 2020). Zheng *el al*. also hypothesized that ASTRAL-Pro is somewhat robust on the underlying rooting and tagging heuristic. . We examined both DISCO and DISCO-R with different scoring heuristics. Table 1 shows six different variants that we have explored in this study.

**Table 1:** Different variants of DISCO analyzed in this study, covering various pruning strategies and scoring schemes for duplication and loss events.

| Variant | Pruning Strategy | Subtree to be pruned | Duplication Weight | Loss Weight |
| --- | --- | --- | --- | --- |
| DISCO-1 | DISCO | Smaller Subtree | 1 | 1 |
| DISCO-2 | DISCO | Larger Subtree | 1 | 1 |
| DISCO-3 | DISCO | Random Subtree | 1 | 1 |
| DISCO-4 | DISCO-R | Smaller Subtree | 1 | 1 |
| DISCO-5 | DISCO | Smaller Subtree | 1 | 0.5 |
| DISCO-6 | DISCO-R | Smaller Subtree | 1 | 0.5 |

## 5 Experimental studies

### 5.1 Datasets

#### Simulated dataset

We analyzed a 25-taxa dataset simulated and analyzed by Zhang et al. (2020) in the ASTRAL-Pro study. For the default model, the number of species is 25, and the number of gene trees is 1,000. The duplication rate (*λ*_+_), which is also equal to the loss rate (*λ*_*−*_), is 4.9 *×* 10^*−*10^ . For the default model, gene trees have an ILS level of [60%, 80%]. We performed most of our experiments with varying duplication rates from 0*∼*5*λ*_+_ and loss/duplication ratios from 0 *∼* 1 with all other parameters kept the same as the default. We also performed some experiments with varying duplication rates and ILS levels while keeping the loss rate equal to the duplication rate.

We also performed an experiment on a dataset of 100 taxa simulated by (Molloy and Warnow, 2020) and also used by (Zhang et al., 2020). The duplication rate was varied as *{*0, 1, 2, 5*} ×* 10^*−*10^, and the loss rate was kept the same as the duplication rate. The ILS was controlled by varying the haploid population size as *{*1, 5*} ×* 10^7^, resulting in a much lower ILS level, 2% and 12%, respectively. Gene trees were estimated considering sequence lengths of 25, 50, 100, and 250 base pairs. and We only considered the model conditions with 500 genes.

#### Empirical dataset

We also tested our methods on two popular biological datasets. First, we use the transcriptome dataset of land plant species (Plants83), which was first analyzed by Wickett et al. (2014). We also reanalyze a dataset of 16 yeast species (Fungi16) with 7,280 multicopy gene trees from Butler et al. (2009) and a dataset of 60 fungal species (Fungi60) with 5659 multicopy gene trees from Morel et al. (2022).

### 5.2 Methods compared

We evaluated different variants of DISCO paired with ASTRAL and wQFM, two leading species tree estimation methods. We also compared wQFM-DISCO (wQFM paired with DISCO) with ASTRAL-Pro, ASTRAL-DISCO, DupTree, and MulRF.

### 5.3 Measurements

We compared the estimated trees (on simulated datasets) with the model species tree using normalized Robinson-Foulds (RF) distance (Robinson and Foulds, 1981) to measure the tree error. The RF distance between two trees is the sum of the bipartitions (splits) induced by one tree but not by the other, and vice versa. All the trees estimated in this study are binary and so False Positive (FP), and False Negative (FN) and RF rates are identical. For the biological dataset, we compared the estimated species trees to the scientific literature. We assessed support values in the estimated trees using local posterior probabilities (Sayyari and Mirarab, 2016) computed by ASTRAL. The local posterior probabilities are computed based on a transformation of normalized quartet scores (the percentage of quartets in the gene trees that agree with a branch in the estimated species tree). We analyzed multiple replicates of data for various model conditions and performed two-sided Wilcoxon signed-rank test (Wilcoxon, 1945). We set the threshold *α* = 0.05 to measure the statistical significance of the differences between two methods.

## 6 Results and discussion

### 6.1 Experiment 1: evaluation of DISCO variants

In Experiment 1, we have two separate experiments to assess: 1(a) the impact of different pruning strategies, and 1(b) the impact of different rooting and tagging strategies.

#### 6.1.1 Experiment 1(a): Effects of choosing the subtree to be pruned

We examined how the performance varies by pruning the smaller (DISCO-1), larger (DISCO-2), or one of them at random (DISCO-3). Originally, DISCO prunes the smaller subtree at a duplication node, but no justification was provided for their choice (Willson et al., 2021). We compared these three methods both with ASTRAL and wQFM with varying duplication rates and loss/duplication ratios (see Figure 3). When the duplication rate is 0, the methods do not differ since the gene trees are single-copy and hence there is no pruning involved. Otherwise, among the 15 other model conditions, we see that ASTRAL-DISCO-1 performs better than both ASTRAL-DISCO-2 and ASTRAL-DISCO-3 on 12 model conditions with estimated gene trees with gene length of 100 bp, on 12 model conditions with estimated gene trees with gene length of 500 bp, and on 11 model conditions with true gene trees. Similarly, wQFM-DISCO-1 showed superior performance compared to wQFM-DISCO-2 and wQFM-DISCO-3 on 11, 11, and 12 model conditions, respectively, across these three types of gene trees. In each case, the difference is statistically significant (*p <<* 0.01). ASTRAL and wQFM paired with DISCO-2 (i.e., pruning the larger trees) produced the worst results in most of the cases. Relative performances of different variants of DISCO paired with either ASTRAL or wQFM are similar. Thus, we experimentally conclude that pruning the smaller subtree is the better strategy. Pruning the smaller subtree retains more speciation-driven quartets, which could be a plausible explanation for our findings. Therefore, in subsequent experiments, we only prune the smaller subtrees as in the original DISCO algorithm.

**Figure 3:**
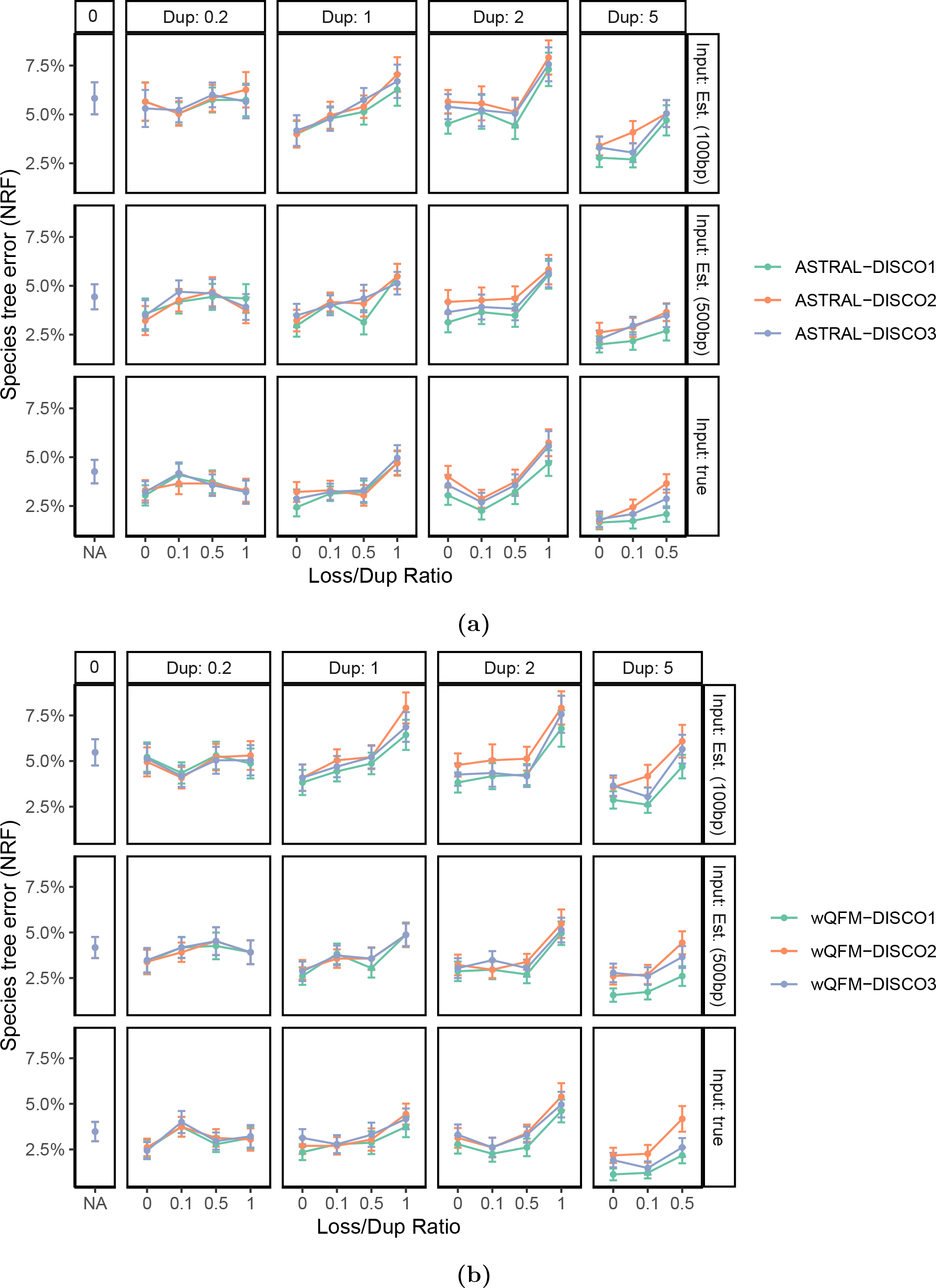
Comparison of variants of DISCO with different pruning techniques, paired with both ASTRAL and wQFM, on the S25 dataset with 1000 gene trees. We show the average RF rates with standard error bars over 50 replicates. We vary the duplication rate (box columns) and the loss rate (x-axis; the ratio of the loss rate to duplication rate). Results for both true gene trees and estimated gene trees from 100 and 500 bp alignments are presented. (a) Results for ASTRAL, paired with different DISCO variants. (b) Results for wQFM paired with DISCO variants.

#### 6.1.2 Experiment 1(b): Effects of different weighting schemes in the tagging technique of ASTRAL-Pro and DISCO

The tagging heuristic used by ASTRAL-Pro (and also by DISCO) equally weights each duplication and loss event. We examined the effects of assigning the weight of a duplication event two times that of a loss event. We also experimented with DISCO-R, the pruning technique we introduced in Section 4. We compared these methods (DISCO-1,4,5,6) paired with both ASTRAL and wQFM under various model conditions (see Figure 4). The relative ranks of the methods are shown in Table 2. Figure 4 and Table 2 suggest that regardless of the choice of species tree estimation methods (i.e., ASTRAL or wQFM), DISCO-R with two different weighting schemes (i.e., DISCO-4 and DISCO-6) are frequently better than original DISCO variants (DISCO-1 and DISCO-5) and the improvements of DISCO-R variants over the original DISCO variants are sometimes statistically significant (*p <* 0.05). However, on true gene trees, all DISCO and DISCO-R variants achieved comparable accuracy, with DISCO variants having a slight advantage over DISCO-R variants on some of the model conditions. In nearly every model condition, DISCO-4 and DISCO-6 methods (paired with ASTRAL or wQFM) had very similar accuracy, with neither reliably having an advantage over the other. Similar observations hold for DISCO-1 and DISCO-5 (i.e,. original DISCO variants with different weighting schemes). This indicates that the rooting and tagging heuristic incorporated by ASTRAL-Pro and also by DISCO is considerably robust to the assignments of weights to duplication and speciation events. These results indicate that our proposed DISCO-R provides more accurate and robust results than the original DISCO algorithm and supports our findings that DISCO-R retains more species-driven quartets than DISCO (as discussed in Sec 4. Therefore, in subsequent experiments, we consider only DISCO-4 (DISCO-R with an equal weighting scheme).

**Table 2:**
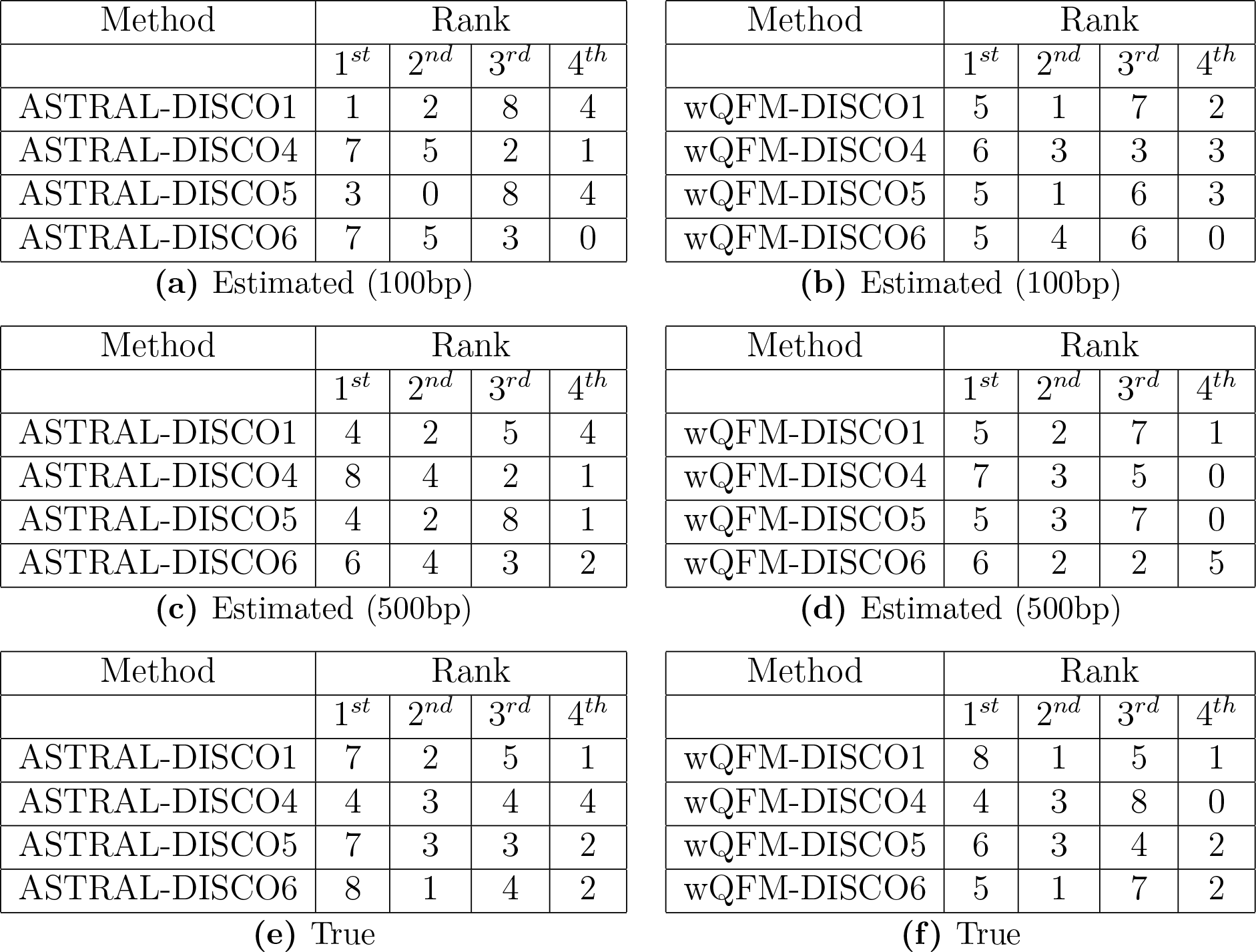
Effects of different weighting schemes in the tagging technique of ASTRAL-Pro and DISCO. We present how the four DISCO variants ranked when combined with both ASTRAL and wQFM are shown in various types of inputs. For example, out of the 15 model conditions with estimated gene trees with 500 bp genes, wQFM-DISCO4 ranked first seven times, second 3 times, and so on. In the case of a group of records with the same value, the lowest rank in the group was assigned.

**Figure 4:**
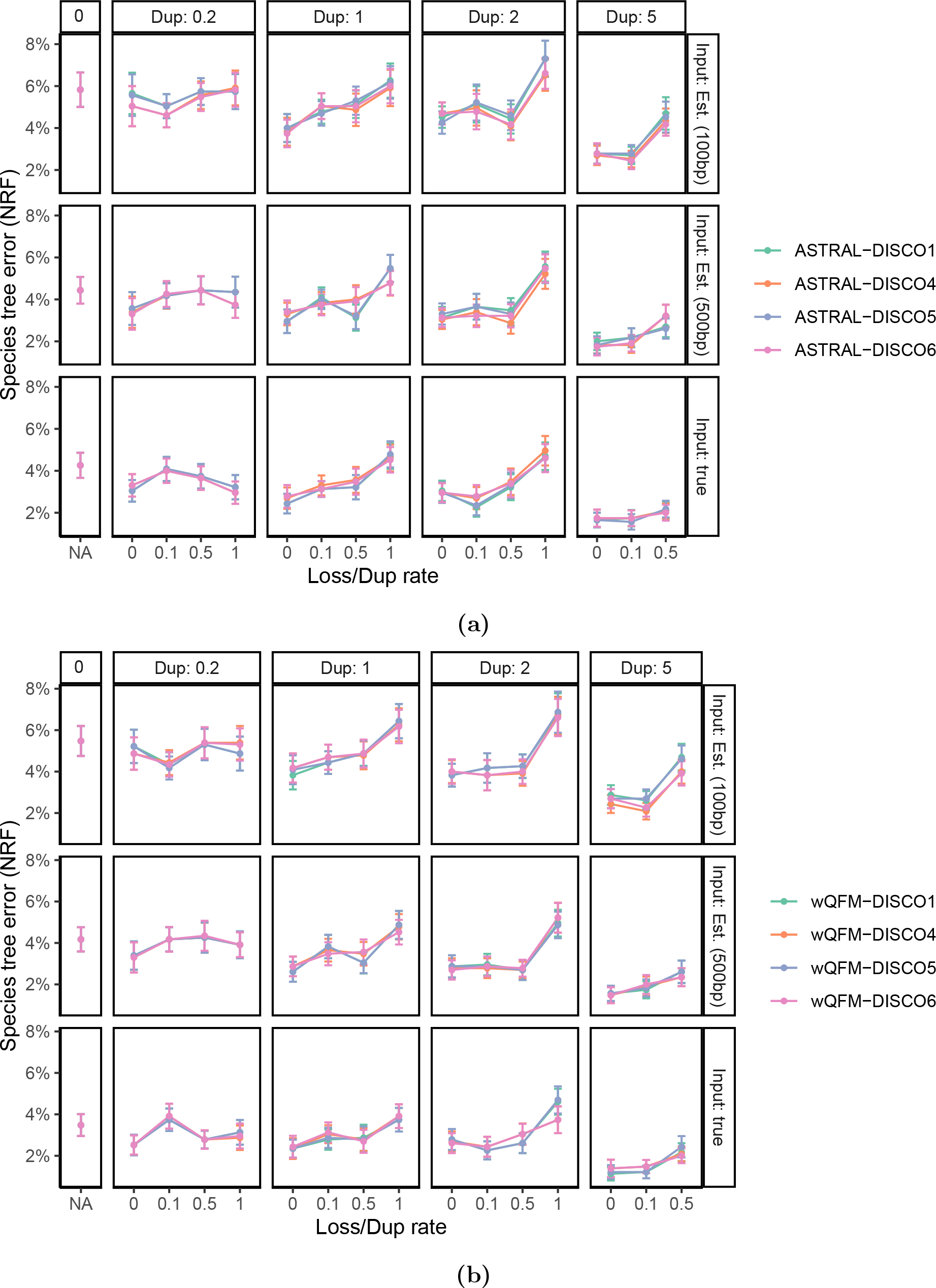
Comparison of variants of DISCO (two different weighting schemes with two different pruning techniques, producing four DISCO variants), paired with both ASTRAL and wQFM, on the S25 dataset with 1000 gene trees. We show the average RF rates with standard error bars over 50 replicates. We vary the duplication rate (box columns) and the loss rate (x-axis; the ratio of the loss rate to duplication rate). Results for both true gene trees and estimated gene trees from 100 and 500 bp alignments are presented. (a) Results for ASTRAL, paired with different DISCO variants. (b) Results for wQFM paired with DISCO variants.

We also observe that in each of the 48 model conditions and 6 variants of DISCO discussed, wQFM paired with any certain variant performed better than ASTRAL paired with that variant in 243 out of 288 times and this improvement is highly statistically significant (*p <<* 0.01). This finding supports the previously obtained result that wQFM is experimentally superior to ASTRAL (Mahbub et al., 2021, 2022).

#### 6.1.3 Experiment 2: Comparisons of best existing summary methods

In this experiment, we compared the best existing method ASTRAL-Pro with wQFM and ASTRAL paired with the best DISCO variant (DISCO-4) identified in our previous experiments. We also included DupTree and MulRF in our comparisons.

##### Varying duplication and loss ratios

First, we do the experiments on the same dataset as analyzed in Experiment 1, where duplication rate and loss/duplication ratio were varied with ILS level fixed at 70%. On true gene trees, in 7 of the 16 model conditions of varying duplication and loss rates, DupTree turns out to be the best method. However, in 7 other conditions, DupTree performed worse than all other methods, except MulRF. However, wQFM-DISCO-4 performed consistently well and it was better than other methods on average. The difference is statistically significant for all methods (*p <* 0.05), except ASTRAL-Pro. MulRF has notably higher errors (*p <<* 0.01) than the other methods across all model conditions.

On the estimated gene trees, the accuracy of DupTree dramatically decreased compared to its performance on the true gene trees. MulRF remained the worst method. wQFM-DISCO-4, ASTRAL-Pro, and ASTRAL-DISCO-4 remained much more accurate than DupTree and MulRF. Remarkably, wQFM-DISCO-4 performs better than ASTRAL-Pro and ASTRAL-DISCO-4 across most of the test conditions. The superior performance of wQFM-DISCO-4 is statistically significant (*p <* 0.05) for estimated gene trees with both 100*bp* and 500*bp* genes.

Of the 16 different combinations of duplication and loss rates under consideration, wQFM-DISCO-4 turned out to be the best method 10, 10, 5 times and the second best method 4, 5, 8 times for estimated trees with 100bp genes, estimated trees with 500bp genes, and true gene trees respectively. On the other hand, ASTRAL-Pro only came out to be the best method 4, 4, 4 times and the second best method 4, 5, 7 times for estimated trees with 100bp genes, estimated trees with 500bp genes, and true gene trees, respectively. In all three types of input, wQFM-DISCO-4 is better on average than the other four methods in consideration.

##### Varying ILS levels

We then repeat this experiment in the case where ILS level also varies with the duplication rate, where loss/duplication rate is kept as 1. MulRF performs very well in conditions with no ILS, but as we increase the level of ILS, the performance of MulRF degrades drastically. wQFM-DISCO-4, ASTRAL-Pro, and ASTRAL-DISCO-4 are relatively robust and much more tolerant of the level of ILS. Remarkably, similar to previous experiments, wQFM-DISCO-4 was the best method. Of the 11 model conditions under consideration, wQFM-DISCO-4 turned out to be the best method 9, 6, 6 times and the second best method 2, 2, 2 times for estimated trees with 100bp genes, estimated trees with 500bp genes, and true gene trees. wQFM-DISCO-4 is either significantly better (*p <* 0.05) than the other four methods, or there is no statistically significant difference between them.

Overall, under a wide range of model conditions with varying duplication/loss and ILS levels as well as varying gene tree estimation errors, wQFM paired with DISCO-4 obtained the best performance. Out of a total 1650 test cases, it performed as good as or better than ASTRAL-Pro in 1503 and the improvement is statistically significant (*p <* 0.05).

##### Varying parameters on the 100 taxa dataset

We then examined the effect of varying gene tree estimation error (controlled by the sequence length), ILS level (controlled by haploid population size), and duplication rate (with duplication/loss ratio kept to 1) in the 100 taxa dataset described in Subsection 5.1. ASTRAL-Pro was reported to be the superior method on this dataset (Zhang et al., 2020), especially when compared to DupTree and MulRF. Consequentially, we report DupTree and MulRF performance in our analysis. As we observe from Fig. 6,wQFM-DISCO-4 turned out to be the best method in 25 out of the 40 model conditions and the second-best method in 8 out of the 15 remaining conditions.

**Figure 5:**
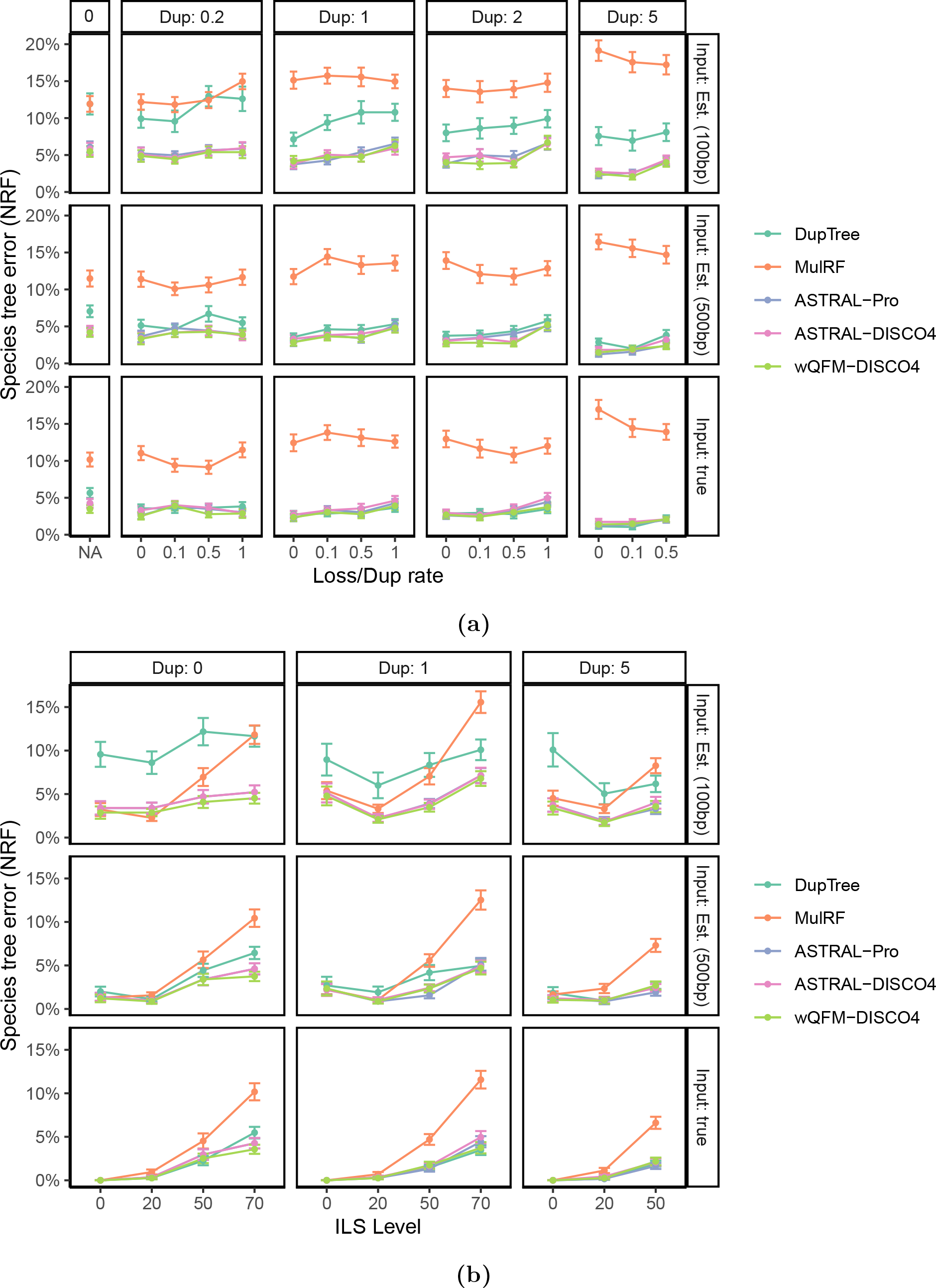
Comparison of wQFM-DISCO-4, ASTRAL-DISCO-4 and some other exisiting methods on the S25 dataset with 1000 gene trees. We show the average RF rates with standard error bars over 50 replicates. We vary the duplication rate (box columns) and the loss rate (x-axis; the ratio of the loss rate to duplication rate). Results for both true gene trees and estimated gene trees from 100 and 500 bp alignments are presented. (a) Results when the duplication rate and loss/duplication ratio was varied. (b) Results when the duplication ratio and ILS level was varied.

**Figure 6:**
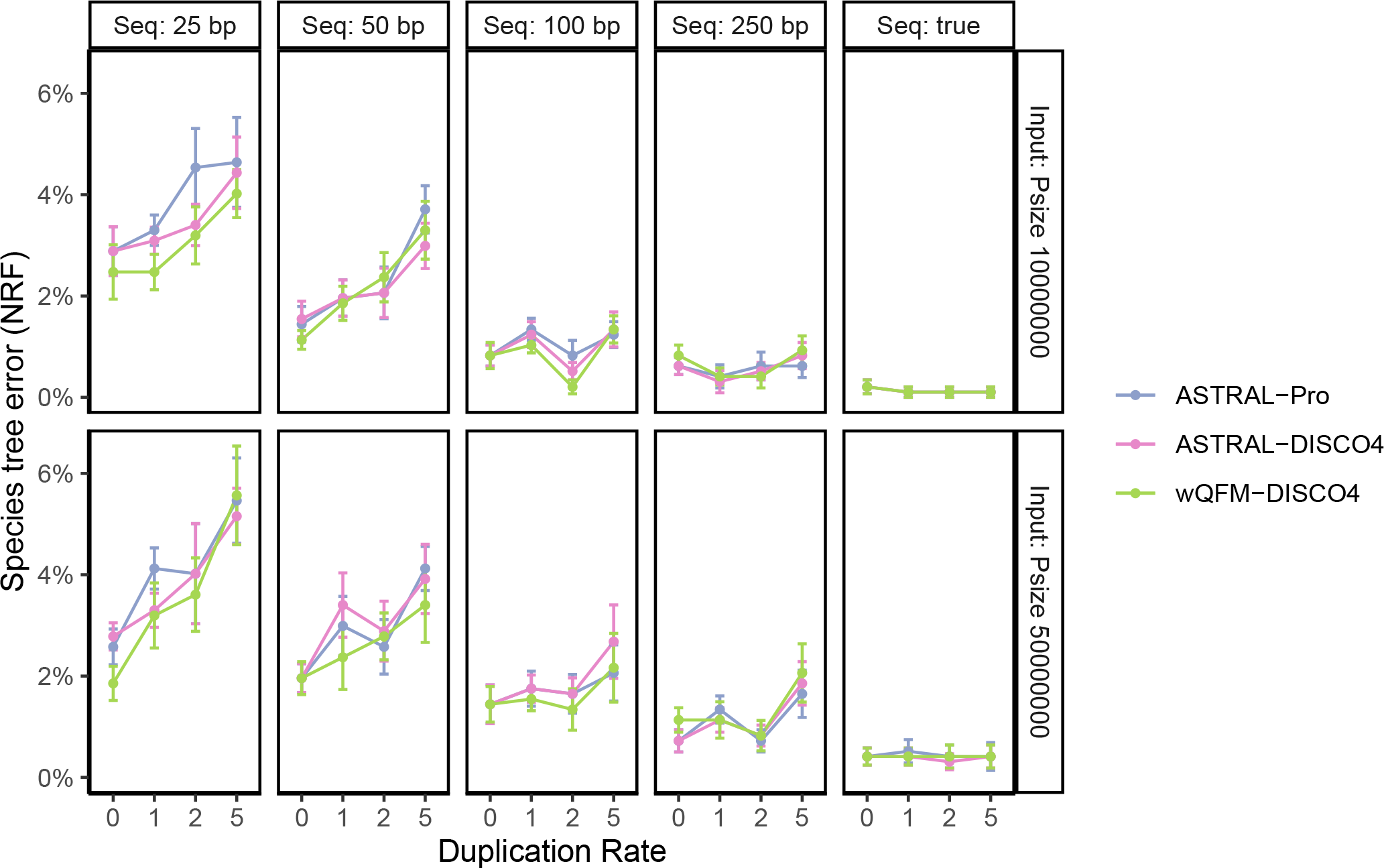
Comparison of wQFM-DISCO-4, ASTRAL-DISCO-4, ASTRAL-Pro on the S100 dataset with 1000 gene trees. We show the average RF rates with standard error bars over 10 replicates. We vary the duplication rate (box columns) and the loss rate (x-axis; the ratio of the loss rate to duplication rate). Results for both true gene trees and estimated gene trees from 25, 50, 100 and 250 bp alignments are presented. The effective haploid population size is also varied.

Overall, in 29 model conditions, wQFM-DISCO-4 performs as good as ASTRAL-Pro or better, and the improvement of wQFM-DISCO-4 over ASTRAL is statistically significant (*p <<* 0.01).

Thus, wQFM is not only superior to ASTRAL on single-copy gene trees (as evident from previous studies (Mahbub et al., 2021, 2022)), wQFM paired with DISCO-4 is more accurate and robust than ASTRAL-Pro under a wide variety of challenging model conditions.

### 6.2 Results on Biological Dataset

#### 6.2.1 Plant (1KP) dataset

We reanalyzed the transcriptome dataset consisting of 103 plant species, which was originally analyzed by Wickett et al. (2014) (Wickett et al., 2014) using 424 single-copy gene trees with the ASTRAL method. The original study inferred 9,683 multicopy gene trees as well, containing up to 2,395 leaves, for 80 out of the 103 species, and three additional genomes, making a total of 83 genomes. However, these multicopy gene trees were not analyzed due to a lack of suitable methods. Later, ASTRAL-Pro analyzed these multicopy gene trees. We reanalyzed this dataset using wQFM paired with both DISCO and DISCO-4. DISCO and DISCO-4 decompose these multicopy gene trees into 55,297 single-copy gene trees.

wQFM-DISCO-4 returned a tree that is highly congruent with the ASTRAL-Pro tree, differing only on three branches with low support (Fig. 7. wQFM-DISCO-4 and ASTRAL-Pro trees differ from the ASTRAL tree based on single-copy gene trees reported by Wickett et al. (2014) in two and five branches, respectively. Both wQFM-DISCO-4 and ASTRAL-Pro exhibit a significant difference from ASTRAL in that they both strongly support the GnePine hypothesis, where Gnetales is sister to Pinaceae, nested within the Coniferales. ASTRAL, on the other hand, recovered the Gnetifier hypothesis (i.e., Gnetales is sister to Coniferales as a whole). ASTRAL-Pro differs from both the ASTRAL and the concatenated analysis using super matirix in the placement of Yucca. In contrast, the placement of Yucca in the wQFM-DISCO-4-estimated tree aligns with ASTRAL and the concatenated analysis. Similarly, ASTRAL-Pro, unlike wQFM-DISCO-4, differs from both ASTRAL and concatenated analysis in the relative position of *Coleochaetale* and *Chara*. ASTRAL-Pro placed Chara and embryophytes as sisters which was rejected by most of the analyses in Wicket et al (Wickett et al., 2014) (see Fig. 4 in Wicket et al). Another discordance with low support is related to the placements of Rosmarinus and Ipomoea, where ASTRAL-Pro, unlike ASTRAL, concatenated analysis and wQFM-DISCO-4, grouped them together as sisters.

**Figure 7:**
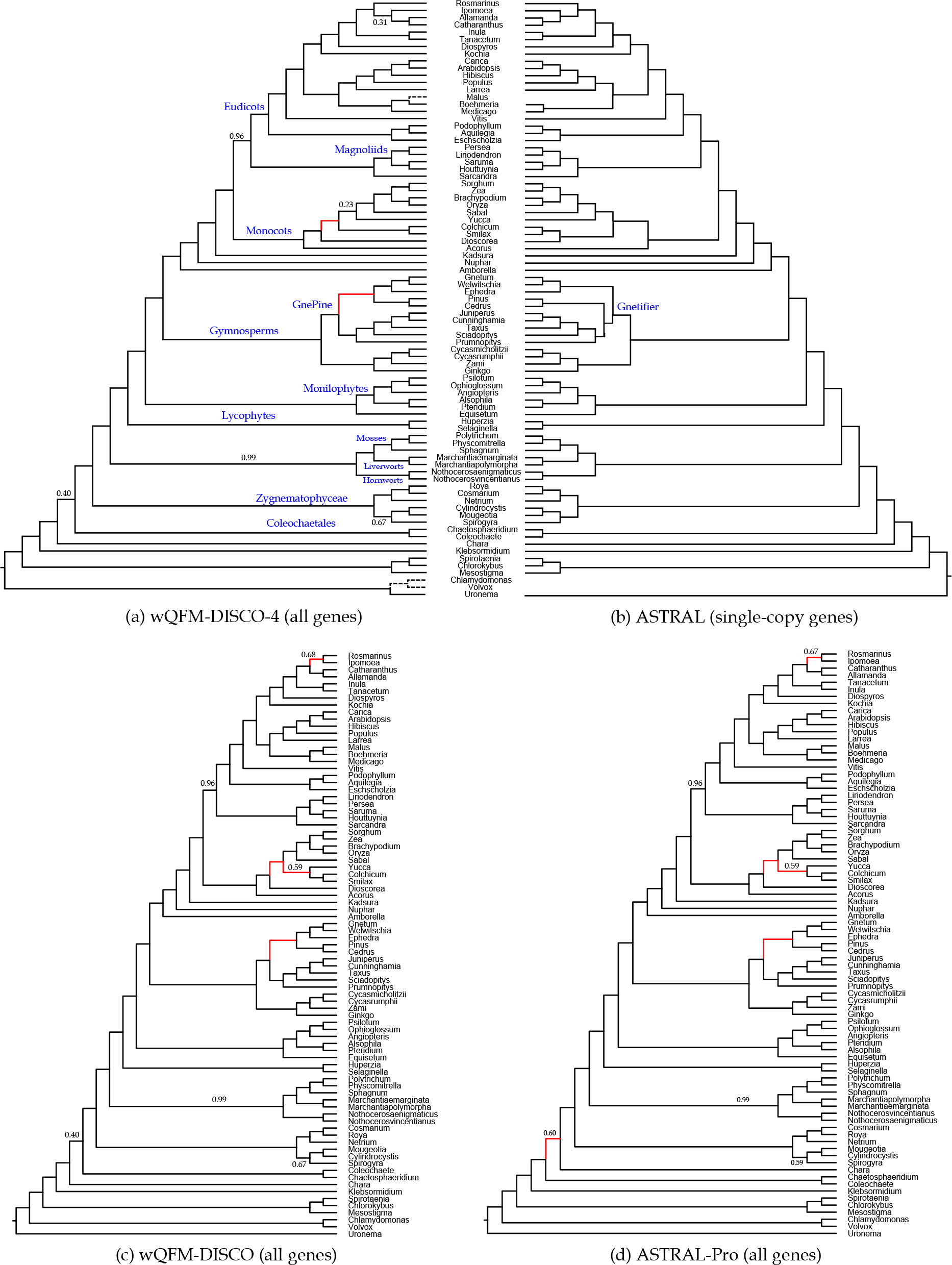
Analysis of the Plants83 dataset. (a) the tree obtained by wQFM-DISCO-4 using all gene trees. Three genomes (indicated by dashed lines) were part of the multicopy dataset but were absent from the single-copy data. (b) the tree reported in (Wickett et al., 2014), estimated by ASTRAL using only the single-copy gene trees. The single-copy tree includes 23 species that were not in the multicopy data and have been removed from the species tree. (c)-(d) trees obtained by wQFM-DISCO and ASTRAL-Pro using multicopy gene trees, respectively. Branch support (BS) values were computed using quartet-based local posterior probabilities. All BS values are 100% except where noted.

Comparing wQFM-DISCO-4 and wQFM-DISCO, we found differences in two branches: 1) the placement of Yucca, and 2) the sister relationships of Rosmarinus and Ipomoea. In relation to these two differences, wQFM-DISCO is aligned with ASTRAL-Pro, while wQFM-DISCO-4 is aligned with the original ASTRAL and concatenated analysis. wQFM-DISCO tree, which exhibits discordance with the ASTRAL tree on four branches, is highly aligned with the ASTRAL-Pro tree, with only one difference pertaining to the placement of chara. Thus, all the methods inferred reasonable species trees, differing from each other in two to five branches (often with low support), where the ground truth is subject to debate.

#### 6.2.2 Fungal dataset

We analyzed two fungal datasets: 1) a data set containing 16 yeast species with 7,280 multicopy gene families available from Butler et al. (2009), and 2) a 60-species dataset from Morel et al. (2022).

##### 16-species yeast dataset

Butler et al. (2009) analyzed only single copy gene trees based on 706 one-to-one orthologs (Butler et al., 2009). We re-analyzed the data considering all the 7,280 multicopy gene trees (see Fig. 8(b). wQFM-DISCO, wQFM-DISCO-4 and ASTRAL-Pro returned an identical tree, which differs from the tree reported in the original study on only one branch, introducing discordance in the relative position of *Saccharomyces castellii* and *Candida glabrata*. The original study placed *Saccharomyces castellii* as sister to *Candida glabrata* and the *Saccharomyces* group which was imposed by a constraint during the ML search. ASTRAL-Pro, wQFM-DISCO and wQFM-DISCO-4 trees placed *Candida glabrata* as sister to the *Saccharomyces* group, which is aligned with the unconstrained ML search in the concatenated analysis and the tree reported by Salichors and Rokas (Salichos and Rokas, 2013).

**Figure 8:**
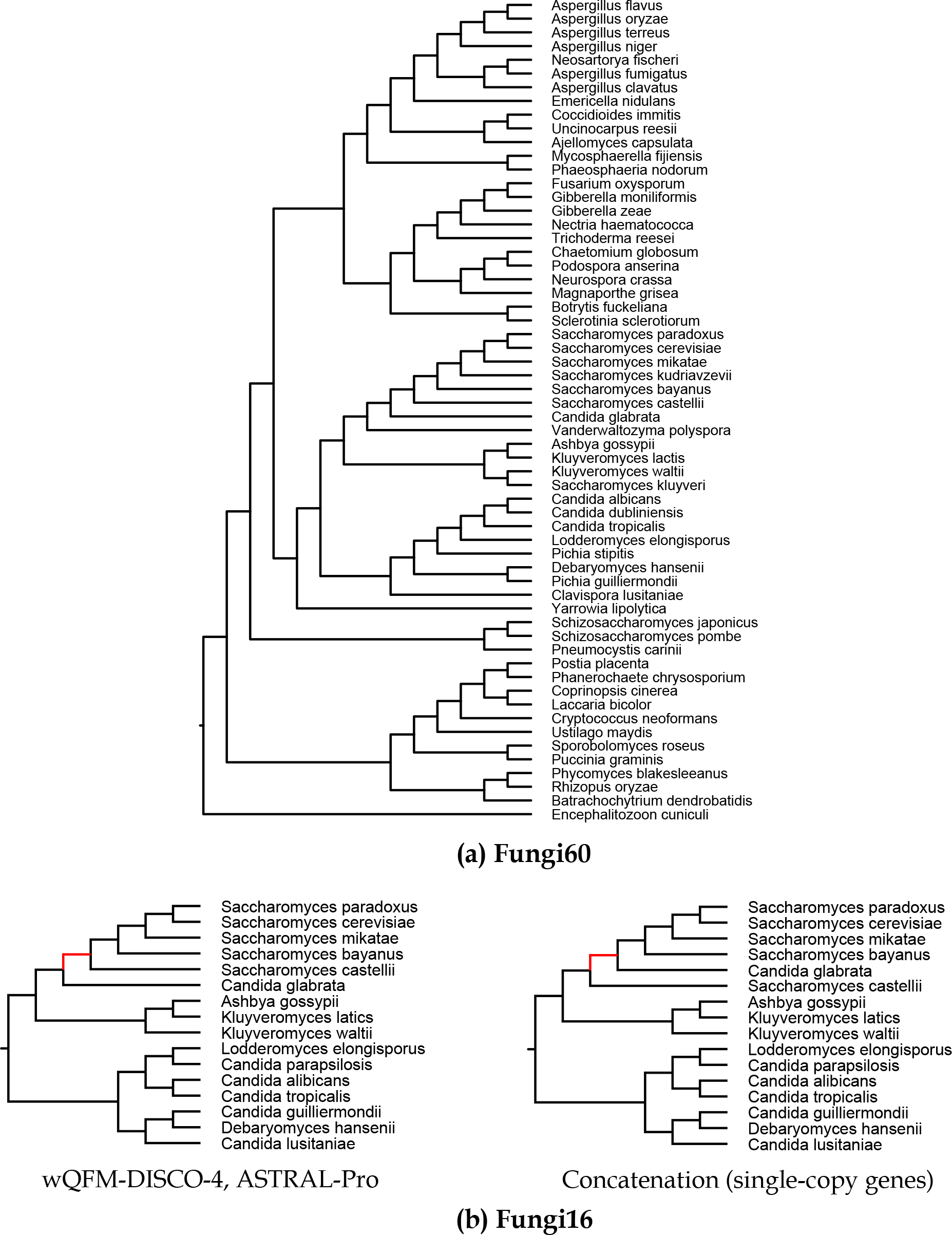
Analysis of the Fungi16 and Fungi60 datasets. (a) Fungi60 dataset: the species tree returned by both wQFM-DISCO-4 and ASTRAL-Pro using all genes. (b) Fungi16 dataset. Left: the species tree returned by both wQFM-DISCO-4 and ASTRAL-pro. Right: the tree obtained by concatenation of 706 single-copy genes with the red branch enforced as a constraint (Butler et al., 2009).

##### 60-species fungal dataset

Morel et al. (2022) extracted a fungal dataset of 60 species from PhylomeDB (Huerta-Cepas et al., 2014). In their study, they found that ASTRAL-Pro, SpeciesRAX and FastMulRFS infer the same tree. In our study, we found that wQFM-DISCO-4 infers the same tree as well, which agrees with the species tree obtained by concatenation (Marcet-Houben and Gabaldón, 2009) (see Fig. 8(a). However, this tree contradicts the relationship established in the literature regarding one specific branch, related to the positioning of the clade comprising Chytridiomycota (*Batrachochytrium dendrobatidis*) and Zygomycota (*Rhizopus oryzae* and *Phycomyces blakesleeanus*). Morel et al. (2022) discussed the biological phenomenon of this split in detail.

### 6.3 Running time

The running time of wQFM is largely dominated by its Ω(|*leaf* |4) quartet generation step from gene trees. When the only cause of gene tree discordance is ILS, the number of leaves equal the number of taxa. However, with GDL in consideration, specially in cases where the duplication rate is much larger than the loss rate, the number of leaves are often times many factors larger than the number of taxon. Running DISCO ensures that the quartet generation step can run in Ω(min(|*leaf* |, |*taxon*|)^4^) time. For analyzing the S100 dataset, the quartet generation took 4 hours on average, but wQFM took only 10 minutes minutes on average to infer a species tree from the weighted quartets. ASTRAL-Pro and ASTRAL-DISCO are faster and takes less than 2 minutes on average to infer the species tree from gene trees. On the 1KP plant dataset with 83 species, generating the quartets from gene trees took seven hours and running wQFM on the quartets took only 5 minutes. ASTRAL-Pro took 39 minutes to infer a species tree. The superior accuracy produced by wQFM further motivates us to focus on how to improve its running time. Especially, if wQFM could be run without explicitly enumerating all quartets from input gene trees (which is the most time-consuming step), it would be much more scalable to large datasets.

## 7 Conclusions

We introduced wQFM-DISCO as a remarkably accurate method for estimating species trees in the presence of gene duplication and loss events. The superior performance of wQFM-DISCO over ASTRAL-Pro across diverse model conditions demonstrates its efficacy and advancement in species tree estimation by effectively modeling both orthology and paralogy. Most of the popular summary methods were originally designed to take ILS into account and cannot handle multicopy gene trees resulting from GDL events. While several parsimony-based summary methods such as DynaDup, DupTree, and iGTP have been developed to consider GDL events, they lack statistical significance. Additionally, traditional approaches like combined analysis or concatenation are not applicable to multicopy gene trees. Reasonably accurate co-estimation methods exist (Boussau et al., 2013b; Zhang and Wu, 2017) but are computationally demanding, limiting their application to small datasets. Hence, species tree estimation in the presence of GDL remains challenging.

ASTRAL-Pro and DISCO are recent advancements in this field. ASTRAL-Pro, a recent summary method, is capable of modeling both orthology and paralogy. DISCO, a simple yet valuable strategy, enables the utilization of methods designed for single-copy gene trees on multicopy gene trees. Our study examined various underlying hypotheses of DISCO and established its robustness by exploring different pruning strategies and scoring schemes for duplication and loss events. We proposed a novel pruning strategy in DISCO, referred to as DISCO-R, which enhances its accuracy and robustness by effectively incorporating more speciation-driven quartets. Finally, we proposed wQFM-DISCO as an adaptation of wQFM for estimating species trees despite the presence of paralogy. Extensive evaluations on diverse simulated and real datasets demonstrated that wQFM-DISCO consistently outperforms ASTRAL-Pro and other competing methods in terms of tree accuracy.

While quartet-based methods like ASTRAL, wQFM, and SVDQuartets are highly accurate and widely used, their application to handle GDL events is limited. ASTRAL-Pro was the first known quartet-based method capable of handling multicopy gene trees and was shown to have superior accuracy compared to other competing methods. wQFM was shown to be more accurate than ASTRAL in multiple studies covering a wide range of model conditions including incomplete gene trees. However, wQFM was not suitable for multicopy gene trees. Therefore, the adaptation of wQFM for multicopy gene trees using DISCO and the demonstrated excellent performance of wQFM-DISCO represent significant advancements in the field.

wQFM, despite being more accurate than ASTRAL, has not gained widespread popularity as a species tree estimation method, perhaps due to its limited scalability when dealing with large datasets containing hundreds of taxa. The presence of gene duplication leads to a substantial increase in the number of leaves, posing an additional challenge to the applicability of wQFM. Importantly, however, the decomposition of multicopy gene trees into single-copy trees using DISCO results in a reduction of leaf numbers, making them more suitable for analysis with wQFM. Thus, wQFM-DISCO is sufficiently fast to analyze the datasets from many important phylogenomic studies (Morel et al., 2022; Wickett et al., 2014). Therefore, wQFM paired with DISCO represents a highly accurate and scalable method to account for both orthology and paralogy, which we believe should be considered as a potential tool to estimate species trees in the presence of ILS and GDL.

This study can be extended in several directions. We analyzed the dataset from the ASTRAL-Pro study covering a wide variety of model conditions. However, future studies need to explore more complex scenarios involving biological processes such as whole genome duplication events, hemiplasy of duplication and loss events, horizontal gene transfer, and others. Although we proved ASTRAL-DISCO to be statistically consistent under the unified model of duplication, loss and coalescence (DLCoal), we could not establish any statistical consistency results for DISCO-R variants. We hypothesize that ASTRAL with DISCO-R would also be statistically significant under DLCoal. There have been some other theoretical results on the MQSST problem (Lafond and Scornavacca, 2019). In particular, the problem is shown to be Fixed Parameter Tractable (FPT), and it also admits a PTAS. Future studies can extend such results in the setting of GDL and explore whether DISCO or DISCO-R can provide any provable approximation guarantee of the optimization problem they try to solve.

## Availability

DISCO-R and other variants are freely available in open source form at https://github.com/skhakim/DISCO-variants. All the datasets analyzed in this paper are from previously published studies and are publicly available.

## Competing interests

The authors declare that they have no competing interests.

## Notes

### Competing Interest Statement

The authors have declared no competing interest.

